# Rapid sequence evolution driven by transposable elements at a virulence locus in a fungal wheat pathogen

**DOI:** 10.1101/2021.02.16.431386

**Authors:** Nikhil Kumar Singh, Thomas Badet, Leen Abraham, Daniel Croll

## Abstract

Background: Plant pathogens cause substantial crop losses in agriculture production and threaten food security. Plants evolved the ability to recognize virulence factors and pathogens have repeatedly escaped recognition due rapid evolutionary change at pathogen virulence loci (*i.e.* effector genes). The presence of transposable elements (TEs) in close physical proximity of effector genes can have important consequences for gene regulation and sequence evolution. Species-wide investigations of effector gene loci remain rare hindering our ability to predict pathogen evolvability.

Results: Here, we performed genome-wide association studies (GWAS) on a highly polymorphic mapping population of 120 isolates of *Zymoseptoria tritici*, the most damaging pathogen of wheat in Europe. We identified a major locus underlying significant variation in reproductive success of the pathogen and damage caused on the wheat cultivar Claro. The most strongly associated locus is intergenic and flanked by genes encoding a predicted effector and a serine type protease, respectively. The center of the locus contained a highly dynamic region consisting of multiple families of TEs. Based on a large global collection of assembled genomes, we show that the virulence locus has undergone substantial recent sequence evolution. Large insertion and deletion events generated length variation between the flanking genes by a factor of seven (5-35 kb). The locus showed also strong signatures of genomic defenses against TEs (*i.e.* RIP) contributing to the rapid diversification of the locus.

Conclusions: In conjunction, our work highlights the power of combining GWAS and population-scale genome analyses to investigate major effect loci in pathogens.

## Background

Plant pathogens are a major threat to food security and cause annual losses of 20-30% of global harvest due to the lack of durable control strategies (Strange & Scott, 2005; Chakraborty & Newton, 2011; Savary, 2020). The emergence of new pathogens, the rise of new virulence in resident pathogens or the gain in resistance against chemical control agents create significant challenges (Strange & Scott, 2005; Subbarao *et al*., 2015; McCann, 2020). To design effective disease control strategies, understanding the molecular interaction between plants and pathogens is critical. The virulence of plant pathogens is largely determined by their repertoire of secreted proteins known as effectors (Rovenich *et al*., 2014; Depotter & Doehlemann, 2020). Effectors target a variety of different plant proteins and metabolic pathways to manipulate the immune response and physiological state of the host (Selin *et al*., 2016). Plants evolved a large array of receptors often organized in networks that can directly or indirectly recognize the presence of effectors (Wu *et al*., 2017; van der Burgh & Joosten, 2019; Depotter & Doehlemann, 2020). Detection of effectors triggers a variety of defense responses preventing the spread of pathogens across plant tissues. The identification of resistance genes encoding receptors has provided key tools for the rapid breeding of resistant crop varieties (Vleeshouwers & Oliver, 2014; Lo Presti *et al*., 2015a). The identification of effectors in plant pathogens is challenging due to the large number of genes encoding effector-like proteins. The repertoire sizes of such candidate effectors varies between filamentous pathogens (Białas *et al*., 2018; Depotter & Doehlemann, 2020). The potato light blight pathogen *Phytophthora infestans* has 1249 predicted effector candidates, whereas the white rust pathogen of *Arabidopsis thaliana, Albugo laibachii*, has only 143 predicted effector candidates (Mcgowan & Fitzpatrick, 2017). The frequent birth and death of genes encoding effectors is underpinning at least part of the variation in candidate effector repertoires among species and underlies also variation within the same species (Fouché *et al*., 2018). Identifying functional effectors providing an advantage for a pathogen on a specific host remains challenging (Selin *et al*., 2016).

Effector gene polymorphism can be a major factor driving host-pathogen interactions (Lo Presti *et al*., 2015b; Fouché *et al*., 2018). The analyses of complete fungal genomes in combination with mapping analyses significantly expanded our knowledge of effectors across major filamentous pathogens. Genome-wide association study (GWAS) and analyses of progeny populations revealed three effectors of the fungal wheat pathogen *Zymoseptoria tritici* (Gohari *et al*., 2015; Zhong *et al*., 2017; Hartmann *et al*., 2017; Stewart *et al*., 2018; Meile *et al*., 2018). The analyses of multiple completely assembled genomes revealed effector genes missing among individual isolates of the species (Plissonneau *et al*., 2016, 2018; Badet *et al*., 2020). Hence, pangenome analyses are crucial to establish the full extent of effector candidates within species (Badet & Croll, 2020). Such effector polymorphism is thought to be at the origin of rapid gains in virulence (Dong *et al*., 2009, 2015; Fouché *et al*., 2018; Asai *et al*., 2018). Breakdown in host resistance can be observed within few years following the deployment of a crop cultivar (Cowger *et al*., 2000; Islam *et al*., 2016; Longya *et al*., 2019; Cowger & Brown, 2019). Effector gene evolution can be driven by the complete deletion of coding sequence, as well as the accumulation of point and frameshift mutations (Rouxel & Balesdent, 2017; Hartmann *et al*., 2017; Fouché *et al*., 2018; Frantzeskakis *et al*., 2020).

The rapid evolution of effector gene sequences is often driven by features of the chromosomal sequence in which the effector genes are embedded. Effector genes can be located on lineage-specific accessory chromosomes (Ma *et al*., 2010; Croll & McDonald, 2012; Manning *et al*., 2013). Such accessory chromosomes are enriched in repetitive sequences (Ma *et al*., 2010). Effector genes located on core chromosomes are often located in the most repetitive regions of the chromosome (Wang *et al*., 2017; Torres *et al*., 2020). The proximity to repetitive regions, in particular transposable elements (TEs), increases the likelihood for sequence rearrangements to occur. The localization of effectors in highly repetitive sub-telomeric regions contributed to rapid virulence evolution of the rice pathogen *Magnaporthe oryzae* (Xue *et al*., 2012; Yoshida *et al*., 2016). The *AVR-Pita* effector gene has been shown to undergo multiple translocations in the genome contributing to the evolution of virulence on specific hosts (Chuma *et al*., 2011). The insertion of a Mg-SINE TE in the effector gene *AvrPi9* led to a loss-of-function mutation enabling *M. oryzae* to escape host resistance (Wu *et al*., 2015). The transposition of TEs can disrupt coding sequences or change the regulation of effector genes (Sánchez-Vallet *et al*., 2018; Meile *et al*., 2018; Fouché *et al*., 2020). Additionally, repetitive sequences can lead to higher mutation rates through a mechanism known as repeat induced point (RIP) mutation (Gladyshev, 2017; Gardiner *et al*., 2020; Wang *et al*., 2020). *Brassica napus* (canola) carrying the *Rlm1* resistance gene suffered a breakdown of resistance against the fungal pathogen *Leptosphaeria maculans* (Van de Wouw *et al*., 2010). The breakdown was associated with a rise in virulence alleles at the *AvrLm1* locus (Van de Wouw *et al*., 2010). Sequence analyses revealed that the gain in virulence was driven by RIP mutations rendering the locus non-functional. Highly similar sequences nearby effector genes can also trigger ectopic recombination and, by this, the deletion or duplication of the effector gene. Consequently, the genomic context of effector genes provides critical information about effector evolvability. Hence, within-species analyses of effector gene diversification and TE dynamics of the surrounding regions have become key tools to retrace the evolution of virulence.

The haploid ascomycete *Zymoseptoria tritici* is one of the most destructive pathogens of wheat leading to yield losses of ∼ 5–30% depending on climatic conditions (Jørgensen *et al*., 2014; Fones & Gurr, 2015). Pathogen populations across the wheat-producing areas of the world harbor significant variation in pathogenicity and genetic diversity (Zhong *et al*., 2017; Hartmann *et al*., 2017; Hartmann & Croll, 2017b; Krishnan *et al*., 2018a; Singh *et al*., 2020). GWAS were successfully used to identify the genetic basis of virulence on two distinct wheat cultivars (Zhong *et al*., 2017; Hartmann *et al*., 2017). In addition, analyses of progeny populations revealed a third effector gene related to a resistance breakdown (Stewart *et al*., 2018; Meile *et al*., 2018). GWAS was also successfully used to map the genetic architecture of a broad range of phenotypic traits related to abiotic stress tolerance (Dutta *et al*., 2020b). TE dynamics are playing a key role in influencing the sequence dynamics at effector gene loci (Hartmann *et al*., 2017; Meile *et al*., 2018; Fouché *et al*., 2020). Gene gain and loss dynamics are accelerated in proximity to TEs (Hartmann & Croll, 2017b). TEs shape also the epigenetic landscape in proximity to effectors (Fouché *et al*., 2020; Meile *et al*., 2020). Phenotypic traits expressed across the life cycle of the pathogen show extensive trade-offs possibly constraining the evolution of virulence (Dutta *et al*., 2020b,a). Identifying additional loci underlying pathogenicity on specific hosts remains a priority as for most wheat resistance (*i.e. Stb*) genes, the matching effector remains unknown (Brown *et al*., 2015).

In this study, we aimed to identify the genetic basis of virulence on the wheat cultivar Claro using GWAS performed on a genetically highly diverse mapping population established from a single wheat field. We analyzed the expression patterns of genes in proximity to the top associated SNP, the presence of TEs and genetic variation at the locus in populations across the world to build a comprehensive picture of sequence dynamics at the newly identified virulence locus.

## Results

### Genome sequencing of a highly polymorphic pathogen field population

In order to build a mapping population for GWAS, we obtained total of 120 isolates of *Z. tritici* collected from a multi-year experimental wheat field in Switzerland planted with 335 wheat cultivars (Karisto *et al*., 2017; Singh *et al*., 2020) (Supplementary Table S1). The isolates were collected from a subset of 10 genetically different winter wheats (7-20 isolates per cultivar) from two different time points during a single growing season. We analyzed whole-genome sequencing datasets of each isolate constituting an average coverage of 21X as previously described (Singh *et al*., 2020). After quality filtering, we obtained 788’313 high-confidence SNPs. We constructed an unrooted phylogenetic network using SplitsTree to visualize the genotypic differentiation within the population (Fig 1A). Compared to the broader field population analyzed previously, our GWAS mapping population contained 10 clonal groups comprising a total of 21 isolates (Singh *et al*., 2020) (Supplementary Table S2). A principal component analysis confirmed the overall genetic differentiation within the population (Supplementary Fig 1). Nearly all genotypes were at similar genetic distances to each other with the exception of five genotypes with significantly larger genetic distances to the main cluster of genotypes (Singh *et al*., 2020) (Supplementary Fig 1). Interestingly, the five isolates were all collected from cultivar CH Combin, which is susceptible to *Z. tritici* (Courvoisier *et al*., 2016).

**Figure 1:**
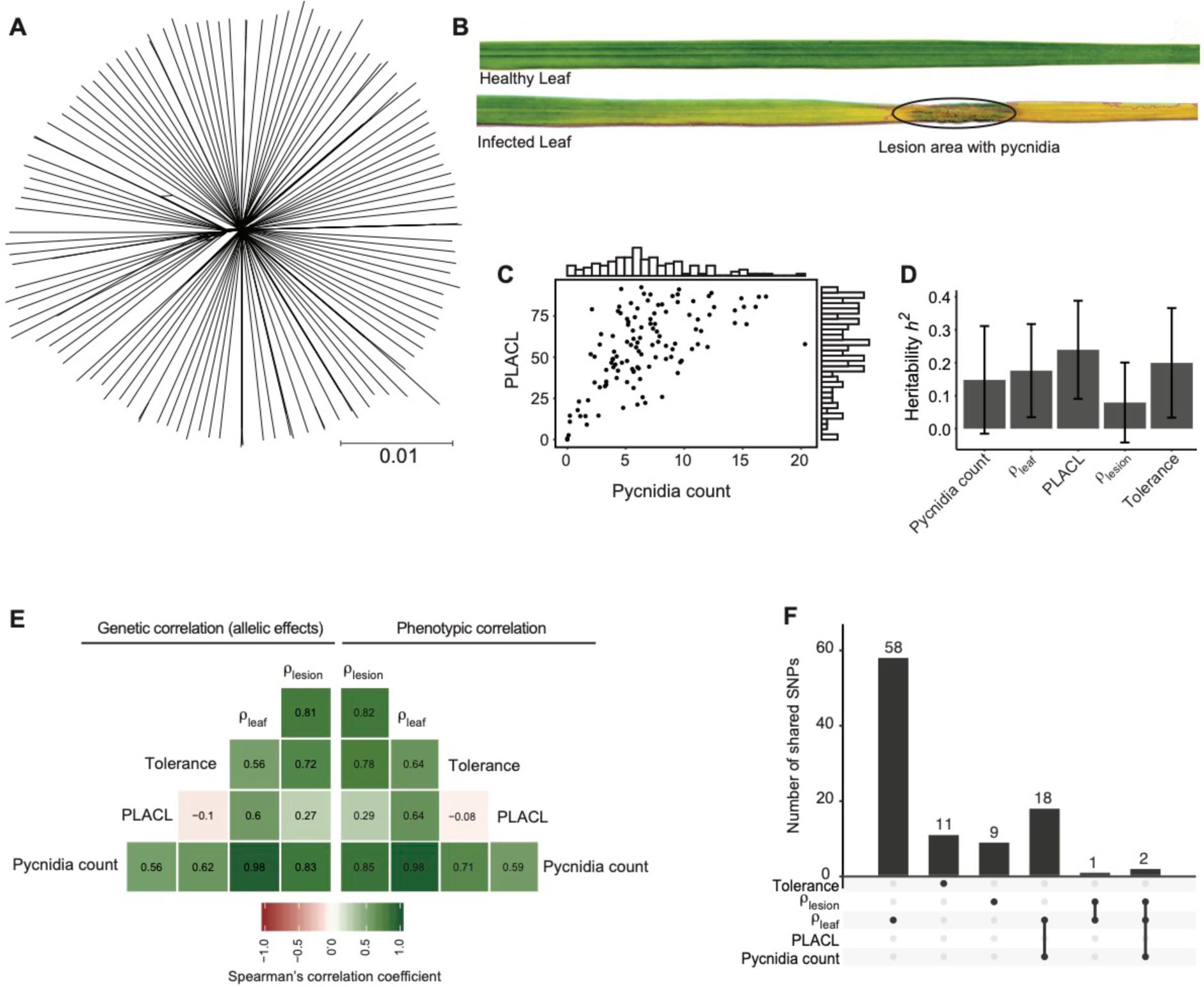
Genetic and phenotypic diversity in a single field population of *Zymoseptoria tritici.* A) Phylogenetic network of 120 isolates constructed using SplitsTree B) Photographs showing the difference between a mock treated and infected leaf. C) Trait distribution of pycnidia counts in lesions and the percentage of leaf area covered by lesion (PLACL). D) SNP based heritability (*h^2^* SNP) of the virulence phenotypes estimated following a GREML approach. Error bars indicate standard errors. E) Mean allelic effect (*i.e.* genetic) correlation and phenotypic correlation coefficients for all measured virulence phenotypes. F) Number of significantly associated SNPs (5% FDR threshold) exclusive to an individual virulence trait or shared among traits.

### Heritability and correlations among pathogenicity traits

We experimentally assessed the expression of pathogenicity traits of each individual isolate on the winter cultivar Claro using a greenhouse assay. The cultivar Claro was among the cultivars used in the multi-year experimental wheat field from which the isolates were sampled from (Karisto *et al*., 2017). The cultivar is widely planted in Switzerland and is generally mildly susceptible to *Z. tritici* (Courvoisier *et al*., 2016). We obtained quantitative data on symptom development from a total of 1’800 inoculated leaves using automated image analysis (Karisto *et al*., 2017). The image analyses pipeline was previously optimized to detect symptoms caused by *Z. tritici* under greenhouse conditions and uses a series of contrast analyses to obtain estimates of the surface covered by symptoms. For each leaf, we recorded the counts of pycnidia (structures containing asexual spores) and the percentage of leaf area covered by lesion (PLACL) (Fig 1B). We considered the pycnidia count as a proxy for reproductive success of the pathogen on the host and PLACL as an indication of host damage due to pathogen infection. From these measurements, we derived three quantitative resistance measures: *ρ*_leaf_ is the pycnidia count per cm^2^ of leaf area, *ρ*_lesion_ is defined as the total number of pycnidia divided by per cm^2^ lesion area, and tolerance is expressed as the pycnidia count divided by PLACL. *ρ*_leaf_ represents the overall reproductive success per area while *ρ*_lesion_ focuses on the reproductive success within the lesion area. Tolerance indicates the ability of the host to tolerate pathogen reproduction while limiting damage by lesions (Mikaberidze & McDonald, 2020). We found that the mean pycnidia count ranged from 0-20 (mean 7, median 6.3) among isolates and PLACP ranged from 2-97 % (mean 56%, median 57.7%) (Fig 1C, Supplementary table S1, Supplementary Fig 2). *ρ*_leaf_ values ranged from 0.04-7.2 (mean 2.4, median 2.15); *ρ*_lesion_ ranged from 0-13.8 (mean 3.6, median 3.3) and tolerance ranged from 0.15-0.3 (mean 0.12, median 0.17) (Supplementary table S1, Supplementary fig 2).

**Figure 2:**
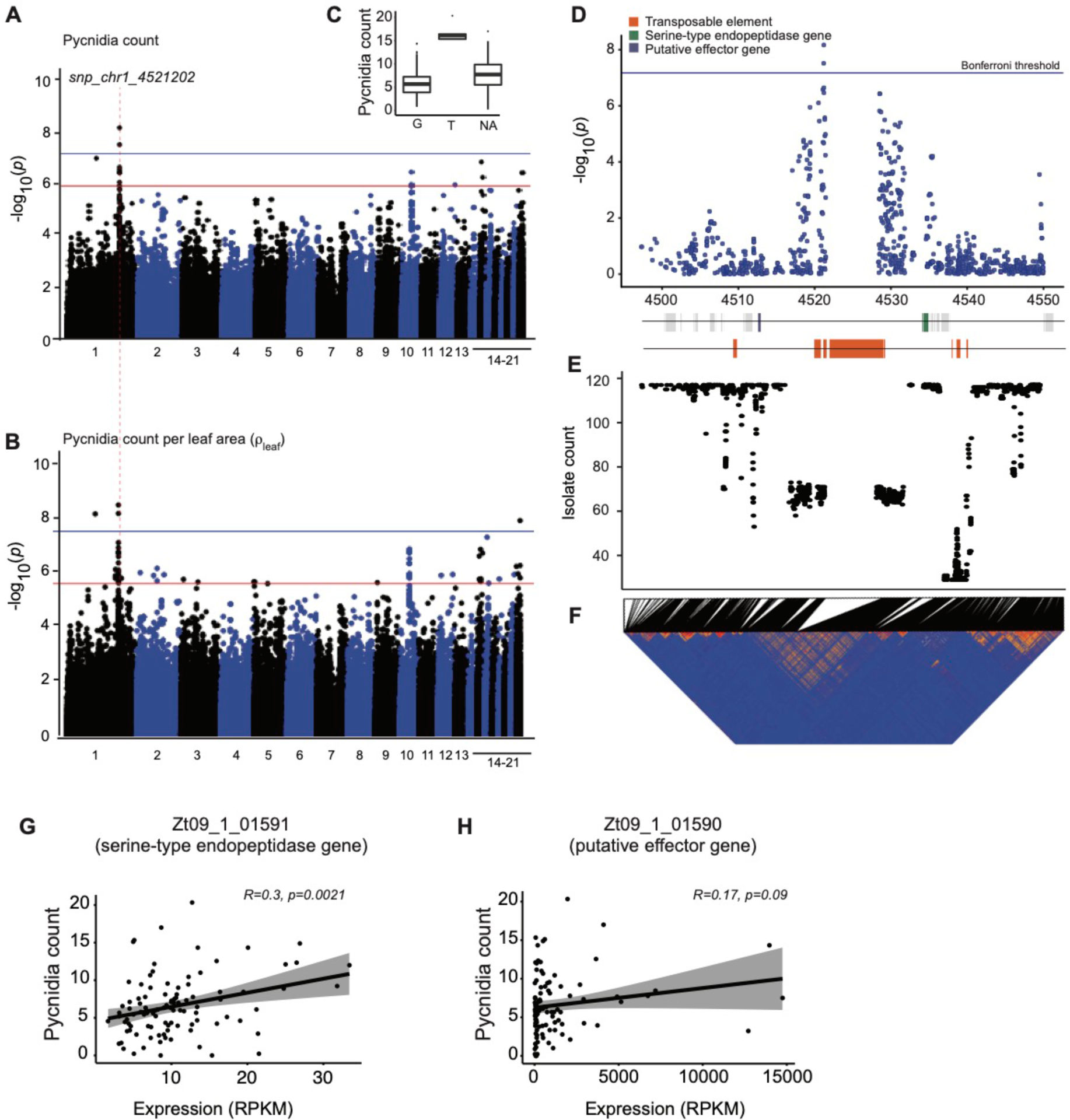
Genome-wide association mapping for virulence. Manhattan plots showing SNP marker association *p-values* for A) pycnidia count and B) ρ_leaf_ (pycnidia count per cm^2^ of leaf area). The blue and red lines indicate the significance thresholds for Bonferroni (*α* = 0.05) and false discovery rate (FDR) at 5%, respectively. The dotted line represents the most significant association on chromosome 1 (*snp_chr1_4521202*). C) Boxplot showing the pycnidia counts of isolates carrying the reference allele G or alternative allele T at the top significant SNP. D) Zommed in Manhattan plot for association *p-values* of SNPs in a ∼25 kb region centered on the top SNP *snp_chr1_452120*. Horizontal lines represent the Bonferroni threshold (*α* = 0.05). E) Genotyping rates of SNPs in the mapping population. F) Linkage disequilibrium *r^2^* heatmap. G-H) Correlation plot of pycnidia count with gene expression of the flanking effector candidate gene (*Zt09_1_01590*) and the serine-type endopeptidase gene (*Zt09_1_01591*).

We estimated SNP-based heritability (*h^2^_snp_*) for each trait using a genomic-relatedness-based restricted maximum-likelihood approach to partition the observed phenotypic variation (Fig 1D). The *h^2^*_snp_ ranged from 0.08 – 0.23 among different phenotypes (Fig 1D). Heritability for pycnidia counts and PLACL was 0.17 (SE = 0.14) and 0.15 (SE = 0.16), respectively. We found the highest *h^2^*_snp_ for *ρ*_leaf_ (0.24, SE=0.15) exceeding *h^2^*_snp_ for *ρ*_lesion_ (0.19, SE=0.16). Pathogenicity-related traits have overlapping genetic architectures leading to phenotypic and genetic correlations (Dutta *et al*., 2020b). To identify potential trade-offs among traits, we analyzed correlations among all pairs of traits (Fig 1E). We found overall positive phenotypic trait correlations except for PLACL and tolerance (*r_p_* = −0.08; Fig 1E). To assess genetic correlations among traits, we performed GWAS on each trait. To avoid *p*-value inflation due to non-random degrees of relatedness among genotypes, we used a mixed linear model that included a kinship matrix. We assessed the allelic effects across all SNPs for all traits to estimate the degree of genetic correlation among trait pairs. We found the genetic correlations (*r_p_*) to vary from −0.1 to 0.98 (Fig 1E). Pycnidia counts and *ρ*_leaf_ showed the highest degree of genetic correlation. PLACL and tolerance showed the lowest degree of genetic correlation. Overall, phenotypic and genetic correlations among pairs of traits were highly similar.

### Major effect locus for pathogen reproduction on the cultivar Claro

We used the GWAS on each trait to identify the most significantly associated SNPs in the genome. We focused on association *p*-values passing the 5% false discovery rate threshold for all the phenotypes except for PLACL where we found no significant associations (Supplementary Fig 3). All significantly associated SNPs for pycnidia count were overlapping with significantly associated SNPs for *ρ*_leaf_ and *ρ*_lesion_ (Fig 1F). The traits *ρ*_leaf,_ *ρ*_lesion_ and tolerance had 58, 9 and 11 associated SNPs, respectively, which were uniquely associated with the specific trait and not overlapping with any other trait (Fig 1F). We then focused our investigation on the most significantly associated SNPs passing the Bonferroni threshold (⍺ = 0.05). We found a single locus on chromosome 1 with significantly associated SNPs for pycnidia count, *ρ*_leaf_ and *ρ*_lesion_ (Fig 2A-B, Supplementary Fig 3C,E). The top SNP (chr1_4521202) showed an association of isolates carrying the non-reference allele T with higher pycnidia production compared isolates with reference allele G (Fig 2C). The non-reference allele was less frequent in the population (10%) and nearly half (48%) of all isolates were not assigned a SNP genotype at the locus.

**Figure 3:**
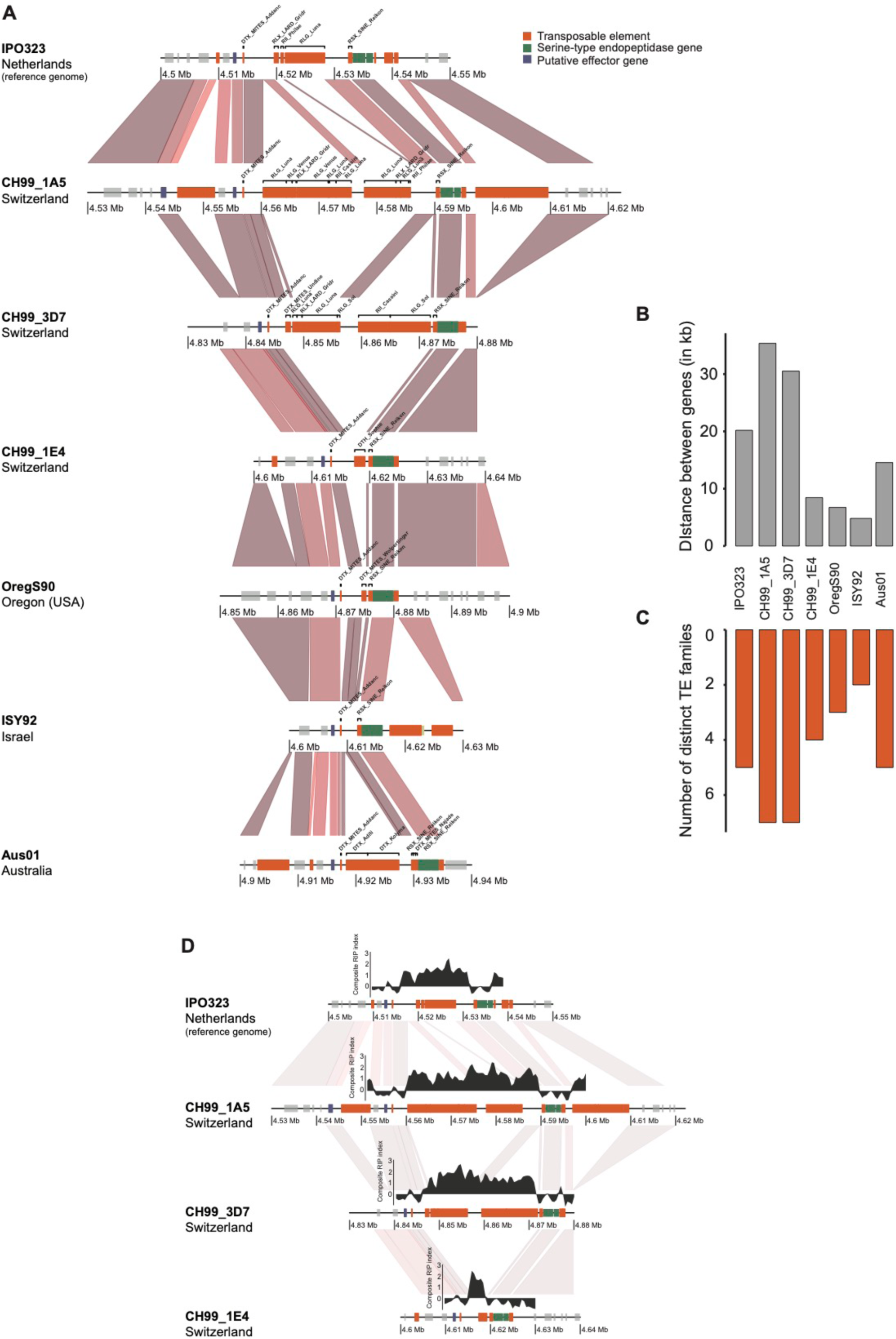
TE content variation at the virulence locus. A) Synteny plot of the top locus analyzed in seven completely assembled genomes. The red gradient segments represent the percentage of sequence identity from BLASTN alignments. Darker colors indicate higher identity. B) Distance variation between the two genes surrounding the top locus (*Zt09_1_01590* and *Zt09_1_01591*). C) The number of different TE families found at least once per isolate at the top locus. D) Repeat induced point (RIP) mutation signatures in the topic locus. The Large RIP Affected Regions (LRARs) composite index was calculated using The RIPper tool (van Wyk et al., 2019).

We analyzed sequence characteristics of the chromosomal region surrounding the top locus. The SNP *chr1_4521202* is located in an intergenic region rich in TEs (Fig 2D-E). The closest identified genes include a gene encoding a putative effector (Zt09_1_01590) and a gene encoding a serine-type endopeptidase (Zt09_1_01591). The genes were at a distance of ∼8 kb and ∼4.5 kb, respectively, from the SNP *chr1_4521202* (Supplementary Table S3). The low genotyping rate at the SNP suggests that segmental deletions are present. The genotyping rate was 58%, which is consistent with the SNP genotyping rate for nearby SNPs (within ∼5 kb; Fig 2E). We recovered no SNPs in the immediate vicinity (at around 4.25 Mb). The genotyping rate increases to close to 100% at a further distance of the top SNP (> 10 kb; Fig 2E. The segmental pattern in the reduced genotyping rate close to the most significant SNP suggests that a substantial fraction of the isolates harbor deletions. We analyzed patterns of linkage disequilibrium among pairs of SNPs (Fig 2F). We found that the decay in linkage disequilibrium generally occurred at short distance (∼1-2 kb) with the exception of the region surrounding the top SNP *chr1_4521202* (Fig 2F). The increased linkage disequilibrium suggests that the physical distance among SNPs in the analyzed isolates is shorter consistent with the detection of deletions.

We analyzed transcription levels of the two closest genes using RNA-seq data generated under culture conditions simulating starvation (minimal medium) for all isolates of the GWAS panel. Both genes were conserved in all the isolates and appear transcriptionally active with variable expression levels among the isolates. The candidate effector gene was transcribed between 12-14’750 RPKM (Fig 2H, Supplementary Fig 4). The serine-type endopeptidase gene showed much lower transcription ranging from 1.6-33.4 RPKM (Fig 2G, Supplementary Fig 4). We found that transcription levels of the gene encoding the endopeptidase was positively correlated with the amount of pycnidia produced (*r* = 0.3, *p* = 0.0021). We found no significant correlation with the expression of the effector candidate gene (Fig 2H).

**Figure 4:**
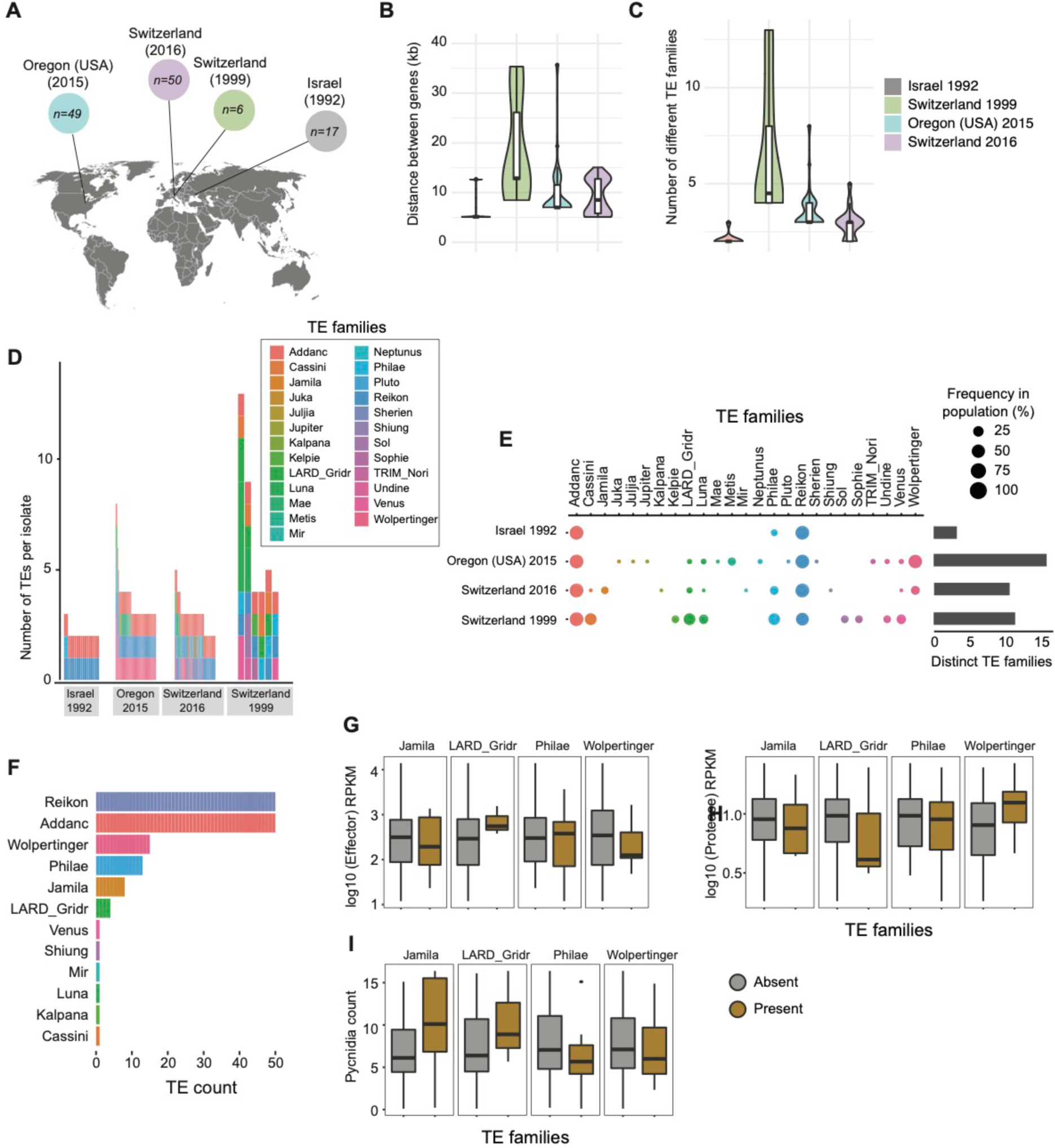
Analysis of transposable element dynamics across continents. A) Analysis of 122 *Zymoseptoria tritici* isolates for which a draft genome assembly produced a scaffold containing both genes *Zt09_1_01590* and *Zt09_1_01591*. B) Boxplot showing variation in the distance between the two genes per population. C) TE content variation of the sequence flanked by the two genes. D) Total TE copies in the sequence flanked by the two genes. E) Frequency of the TE families in the sequence flanked by the two genes as a percentage of the population. F) Frequency of TE families among the isolates from the GWAS mapping population (*n* = 50 with a scaffold spanning both genes) G-H) Boxplots showing the expression of the genes *Zt09_1_01590* and *Zt09_1_01591*, and pycnidia counts for isolates carrying or not specific TEs at the top locus.

### Transposable element dynamics and sequence rearrangements

Given the indications for segmental deletions at the virulence locus, we analyzed multiple completely assembled genomes of the species. We included genomes from isolates from Switzerland, United States, Australia and Israel covering the global distribution range of the pathogen (Badet *et al*., 2020). The locus showed a highly variable content in TEs underlying significant length variation. The distance by the two flanking genes is 20.2 kb in the reference genome IPO323 used for mapping (Fig 3A-B). However, this distance varies from 4.8–35.3 kb between the genes depending on the genome for an average distance of ∼17 kb (Fig 3B). The longest distance between genes was found in the genome of the Swiss strain CH99_1A5 and the shortest distance was found in the genome of the Israeli strain ISY92.

We identified five different TE families in the reference genome IPO323 covering a segment of ∼20 kb (Fig 3C). We detected additional TE families in two of the three genomes from Switzerland (CH99_1A5 and CH99_3D7). The genomes carry multiple copies of a total of seven different TE families. Meanwhile, the two genomes from Israel and the United States showed a reduction in TEs with the region carrying only single copies of two and three different TE families, respectively (Fig 3A-C). The presence of TEs in fungal genomes can trigger RIP mutations. We found consistent signatures of RIP between the two flanking genes but we found no indications for RIP leakage into the flanking genes (Fig 3D, Supplementary Fig 5).

### Transposable element insertion dynamics across populations

The small set of completely assembled genomes provides only a partial view on the sequence rearrangement dynamics within the species. Hence, we generated draft genome assemblies for 432 isolates from previously analyzed field populations in the United States (n = 56 + 97), Switzerland (n = 37 + 185), Israel (n = 30) and Australia (n = 27; Supplementary table S4). The two population from United States and Switzerland were collected at an interval of 25 and 20 years, respectively, from the same field (Supplementary table S4). Illumina sequencing datasets for fungi with compact genomes produce reasonably accurate draft assemblies (Torriani *et al*., 2011; Mohd-Assaad *et al*., 2019; Stauber *et al*., 2020). We used BLASTN (Altschul *et al*., 1990) to locate the two genes *Zt09_1_01590* and *Zt09_1_01591* adjacent to the top SNP across all assemblies. We retained only draft assemblies for which both genes were located on the same scaffold. Hence, these scaffolds provide a contiguous view on the sequences located between the two adjacent genes. With this filtering step, we retained 122 isolates from all four different locations including the United States (n=49), Israel (n=17) and Switzerland (n =6 and 50) (Fig. 4A; Supplementary table S4). The distance between the two genes ranged from 5-35 kb, which is highly consistent with the gene distances observed in the completely assembled genomes (Fig 4B). The isolates from Israel and Switzerland (collection 2016) showed shorter distance ranging from 5-15 kb. The United States population and the older Switzerland population (collection 1999) showed a range of 6.8-35 kb between the genes (Fig 4B).

We annotated the scaffolds matching the top GWAS locus using consensus sequences of known TE families. Overall, the TEs between the two adjacent genes grouped into 11 superfamilies and 25 families (Fig 4D). The two most frequent TEs included both retrotransposons and miniature-inverted repeat transposable elements. We found that genomes from the United States and the earlier Switzerland population (collection 1999) had higher TE copy numbers compared to other genomes from the other populations (2-13 TE copies; Fig 4C). The locus contains overall 16 different TE families in the United States population (Fig 4D-E). The locus contained 3, 11 and 16 different TE families in the Israel, and the Switzerland 1999 and 2016 populations, respectively (Fig 4D-E). The TEs RSX_SINE_Reikon and DTX_MITES_Addanc were found in all analyzed populations population while other TE families were segregated in populations in different proportions (Fig 4E). To test for potential associations of TE presence and pathogenicity traits, we focused on the complete scaffolds retrieved from 50 different isolates of the GWAS population. We found segregating presence-absence polymorphism for the four TE families RII_Philae and RLX_LARD_Gridr (retrotransposons), as well as DTX_MITES_Wolpertinger and DTC_Jamila (DNA transposons; Fig 4F). None of the TE presence-absence polymorphism showed a significant association with the transcription of adjacent genes (effector candidate *Zt09_1_01590*: Student’s *t*-test, *p* > 0.15; serine-type endopeptidase *Zt09_1_01591*: Student’s *t*-test, *p* > 0.05). We also found no significant association with the TE presence-absence polymorphism and any of the pathogenicity traits (Student’s *t*-test, *p* > 0.2).

## Discussion

We used whole-genome sequencing data and association mapping to unravel the genetic architecture of pathogenicity of *Z. tritici* on the wheat cultivar Claro. The identified locus is rich in TEs and is flanked by genes encoding an effector candidate and a serine type protease. We analyzed a worldwide set of populations to analyze sequence variation at the pathogenicity locus. We found significant length variation caused by the insertion of a diverse set of TEs.

Variation in pathogenicity on the wheat cultivar Claro was largely quantitative. We found that heritability was higher for pathogen virulence (damage to host) than pathogen reproduction (production of pycnidia). This is in contrast to analyses of heritability across 12 different wheat cultivars where heritability for pathogen reproduction was typically higher compared to lesion damage (Dutta *et al*., 2020b). However, virulence and reproduction were overall positively correlated in both studies. We also found a high degree of phenotypic and genetic correlation with tolerance (*i.e.* preventing lesion damage despite high reproduction of the pathogen). Using GWAS, we identified several loci significantly associated with different pathogenicity traits. The most significant associations were found for pycnidia counts and *ρ*_leaf_ both related to reproductive success of the pathogen. Pathogen reproduction showed a strong single locus association while host damage (*i.e.* lesions) revealed no single gene effects. The difference between traits may be due to the fact that the genetics underlying host damage is more complex. Lesions are caused by host cell death triggered as a response to pathogen attack (Coll *et al*., 2011; Dickman & de Figueiredo, 2013). Hence, variation in lesion development among isolates could be due to the host’s ability to perceive specific molecules produced only by a subset of the isolates. Furthermore, variation in the pathogen’s ability to spread across tissue and manipulate host immune responses could also lead to variation in overall lesion development. Interestingly, extensive lesion development is not necessarily related to pycnidia production by the pathogen across cultivars (Karisto *et al*., 2018; Mikaberidze & McDonald, 2020). This suggests that despite damage to the leaf, the host immune system can efficiently repress the pathogen from acquiring nutrients to reproduce. In contrast, the strong single locus association for pycnidia production on the cultivar Claro suggests a rather simple genetic architecture. Hence, the action of a single pathogen factor (*e.g.* an effector) may be largely sufficient to determine variation in host exploitation and reproduction.

We identified a highly polymorphic chromosomal locus associated with pathogenicity on the cultivar Claro. The most significant SNPs mapped in an intergenic region flanked by a large cluster of diverse TEs. We found no clear evidence for a coding sequence in immediate proximity of the most significantly associated SNPs. The closest genes encode functions, which may be relevant for host infection though. Serine-type endopeptidases play pivotal roles in nutrient degradation and subsequent assimilation, as well as protection from the host immune system (Muszewska *et al*., 2017). Serine proteases can also help the pathogen to escape the host’s immune system by degrading chitinases targeted at the fungal cell wall (Langner & Göhre, 2016). Furthermore, serine proteases play a role in the nutrient acquisition from plant tissue (Jashni *et al*., 2015) and potentially during the initiation of necrosis (Palma-Guerrero *et al*., 2016). The second gene encodes a putative effector, which is a category of genes showing frequent presence-absence polymorphism within the species (Hartmann & Croll, 2017a; Badet & Croll, 2020). Our analyses of linkage disequilibrium decay suggest that neither of the two adjacent genes play a causal role in pathogenicity on Claro. The most dramatic changes occurred due to the insertion and deletion of TEs next to the most significantly associated SNPs. The TE dynamics diversified the locus to the extent that the distance between the adjacent genes varies by a factor of seven (5-35 kb). The insertion and deletion of TEs can have both an impact on gene regulation by inducing epigenetic silencing or upregulation. Both mechanisms are well established in *Z. tritici* and underlie variation in melanin production, virulence and fungicide resistance (Omrane *et al*., 2017; Meile *et al*., 2018; Krishnan *et al*., 2018b). The locus flanked by the two genes showed strong signatures of RIP. The elevated mutation rates triggered by this genomic defense mechanism against TEs likely contributed to the rapid diversification of the locus. Yet, we could not establish any direct association between the insertion of individual TEs and the expression of pathogenicity. Targeted deletion assays focusing on individual sequence segments may provide experimental evidence for the sequence variation underlying pathogenicity on the wheat cultivar.

### Conclusions

The effects of gene-TE proximity have been studied mainly in animal (Rebollo *et al*., 2012; Cowley & Oakey, 2013) or plant models (Bennetzen & Wang, 2014). Only a handful studies are focused on fungi. Some fungal pathogens have genomes with a clearly compartmentalized architecture described by the two-speed model (Dong *et al*., 2015). The core genome encodes all essential genes while niche- or host-specific genes (*e.g.* effectors) are typically encoded in the repeat-rich genome compartment. Such genome architectures have been identified in *Mycosphaerella fijiensis* (Santana *et al*., 2012), *Cochliobolus heterostrophus* (Santana *et al*., 2014)*, Fusarium* species (Sperschneider *et al*., 2015)*, Leptosphaeria maculans* (Rouxel & Balesdent, 2017) and *Verticillium* species (Faino *et al*., 2016). However, systematic investigation of TEs and co-localizing genes have rarely been extended to the within species level. Our study shows that a combination of genome-wide association mapping, complete and draft genome assemblies can provide a comprehensive insight into the evolutionary dynamics of virulence loci. Hence, even in absence of experimentally validated effectors, the evolutionary trajectory of virulence loci becomes tractable. Our approach should be broadly applicable to many fungal pathogen system.

## Methods

### Field collection and storage

*Z. tritici* isolates were collected from the Field Phenotyping Platform (FIP) site of the ETH Zürich, Switzerland (Eschikon, coordinates 47.449°N, 8.682°E). We analyzed a total of 120 isolates collected during the 2015/2016 growing season from 10 winter wheat cultivars, which are commonly grown in Switzerland (Levy et al., 2017). We analyzed isolates originating from two collection time points over the season (Table S1). Isolates from the first collection (*n* = 62) were collected when wheat plants were in Growth stage (GS) 41 while the second collection (*n* = 58) was performed when the plants were in GS 85 stage. After sampling, spores of each isolate were stored in either 50% glycerol or anhydrous silica gel at −80 °C. Additional information regarding sampling schemes and genetic diversity is available (Singh *et al*., 2020)

### Culture preparation and seedling infection assay

Isolates were revived from glycerol stock by adding 50 µl fungal stock solution to a 50 ml conical flask containing 35 ml liquid YSB (yeast-sucrose broth) medium. The inoculated flasks were incubated in the dark at 18^°^ C and 140-180 rpm on a shaker-incubator. After 8 days of incubation, the cultures were passed through four layers of meshed cheesecloth and washed twice with sterile water to remove media traces. The filtering step also largely eliminated hyphal biomass but retained spores. The Swiss winter wheat cultivar Claro was used for virulence assays (provided by DSP Delley, Inc.). Four seeds were sown in pots with commercial compost soil in triplicates. The pots were frequently in the growth chamber. The plants were grown under controlled conditions as follows: 16/8 hours day/night periods at 18°C throughout the experiment. The growth chamber was maintained at 70% humidity. Plants were grown for three weeks before infection with *Z. tritici*. To initiate infections, washed spores were diluted to 2 x 10^5^ spores/ml in 15 ml of sterile water containing 0.1% TWEEN20. For each isolate, plants from three pots were infected using spray bottles. After spray inoculation, the plants were allowed to dry before sealing them in clear plastic bags to maintain 100% humidity for 48 hours. Plastic bags were removed after 48 hours and conditions were kept as described above.

### Automated image-based evaluation of infection

Twenty-one days post inoculation (dpi), the second leaf of each plant was cut and fixed on a barcoded white paper. Leaves were scanned immediately using a flatbed scanner at 1200 dpi. The scanned images were batch-processed using a macro (Stewart *et al*., 2016; Karisto *et al*., 2017) based on routines implemented in the image analysis software ImageJ (Rasband, W.S., ImageJ; U. S. National Institutes of Health, http://imagej.nih.gov/ij/, 1997–2012). Briefly, the macro recorded the total leaf area, total lesion area, the number of pycnidia, mean size of pycnidia and pycnidia grey value. The percent leaf area covered by lesions (PLACL) was calculated as the ratio of the total lesion area and total leaf area (Karisto *et al*., 2018).

### Whole-genome sequencing, variant calling and RNA-seq analyses

Approximately 100 mg of lyophilized spores were used to extract high-quality genomic DNA using the Qiagen DNeasy Plant Mini Kit according to the manufacturer’s protocol. We sequenced paired-end reads of 100 bp each with an insert size of ∼550 bp on the Illumina HiSeq 4000 platform. Raw reads are available on the NCBI Sequence Read Archive under the BioProject PRJNA596434 (Oggenfuss *et al*., 2020). For RNA sequencing, the same isolates were cultured in a Vogel Minimal N Medium (Vogel, 1956) where ammonium nitrate was replaced with potassium nitrate and ammonium phosphate (Metzenberg, 2003). The medium contained no sucrose and agarose in order to induce hyphal growth. Total RNA was isolated from the filtered mycelium after 10-15 days using the NucleoSpin® RNA Plant and Fungi kit. The RNA concentration and integrity were checked using a Qubit 2.0 Fluorometer and an Agilent 4200 TapeStation System, respectively. Only high-quality RNA (RIN>8) was used to prepare TruSeq stranded mRNA libraries with a 150 bp insert size and sequenced on an Illumina HiSeq 4000 in the single-end mode for 100 bp.

### Sequencing filtering and analysis

We performed sequencing quality checks using FastQC v. 0.11.9. (Andrews S., 2010) and extracted read counts. Sequencing reads were then trimmed for adapter sequences and sequencing quality using Trimmomatic v. 0.39 (Bolger et al., 2014) using the following settings: illuminaclip=TruSeq3-PE.fa:2:30:10, leading=10, trailing=10, sliding-window=5:10 and minlen=50. Trimmed sequencing reads were aligned to the reference genome IPO323 (Goodwin et al., 2011); accessible from https://fungi.ensembl.org/Zymoseptoria_tritici/Info/Index) and the mitochondrial sequence (European Nucleotide Archive EU090238.1) using Bowtie2 v. 2.4.1 (Langmead & Salzberg, 2012). Multi-sample joint variant calling was performed using the HaplotypeCaller and GenotypeGVCF tools of the GATK package v. 4.0.1.2 (McKenna et al., 2010). We retained only SNP variants (excluding indels) and proceeded to hard filtering using the GATK VariantFiltration tool based on the following cutoffs: QD < 5.0; QUAL < 1000.0; MQ < 20.0; −2 > ReadPosRankSum > 2.0; −2 > MQRankSum > 2.0; −2 > BaseQRankSum > 2.0. After filtering for locus level genotyping rate (>50%) and minor allele count (MAC) of 1 using VCFtools v. 0.1.15 (Danecek et al., 2011). Similarly, RNA sequences were checked for quality using FastQC v. 0.11.9. and trimmed with Trimmomatic v0.39 to remove adapter sequences and low-quality reads with parameters: illuminaclip:TruSeq3-SE.fa:2:30:10 leading=3, trailing=3, sliding-window=4:15 and minlen=36. Trimmed sequences were aligned to the reference genome IPO323 using HISAT2 v. 2.1.0 (source?) with the parameter “--RNA-strandedness reverse”.

### Population genetic analyses

Population structure and relatedness among individuals in the mapping population may be a source of *p*-value inflation due to non-random phenotype-genotype associations (Bergelson & Roux, 2010; Korte & Farlow, 2013). To account for this, we analyzed the population structure and genetic relatedness of all isolates by performing a principal component analysis (PCA). We performed and visualized the PCA using the R packages vcfR v. 1.8.0 (Knaus & Grünwald, 2017), adegenet v. 2.1.1 (Jombart & Ahmed, 2011), and ggplot2 v. 3.1.0 (Wickham, 2016). We also generated an unrooted phylogenetic network using SplitsTree v4.14.6 (Huson, 1998). File format conversions were performed using PGDSpider v2.1.1.5 (Lischer & Excoffier, 2012). To identify groups of clonal genotypes, we calculated the pairwise genetic distances between all genotypes using the function “dist.dna” included in the R package ape v. 5.3 (Paradis & Schliep, 2019). Isolate pairs with a pairwise genetic distance below 0.01 were considered as clones for further analyses (see (Singh *et al*., 2020) for more details). The SNP-based heritability (*h^2^*_snp_; equivalent to narrow-sense heritability) for each trait was estimated using the genome-wide complex trait analysis (GCTA) tool v.1.93.0 (Yang *et al*., 2011). The *h^2^*_snp_ was estimated using a genome-based restricted maximum likelihood (GREML) approach using the phenotypic values of each trait and considering the additive effect of all the SNPs represented by the GRM.

### Genome-wide association mapping and linkage disequilibrium analyses

We performed GWAS based on mixed linear models accounting for kinship (MLM K). We first estimated relatedness among genotypes by computing a kinship matrix using the scaled identity by state (IBS) algorithm implemented in TASSEL v. 20201110 (Bradbury *et al*., 2007). We included the kinship matrix as a random effect in the mixed linear models for association mapping using TASSEL. We used the allelic effect output of TASSEL to compute the pairwise genetic correlation (Spearman’s correlation) values using complete observations (use = pairwise.complete.obs) and visualized the values using the *ggcorr* function from the GGally R package (source). Similarly, we used the Spearman’s correlation to compute pairwise phenotypic trait correlations. Association mapping outcomes were visualized using the R package *qqman* v 0.1.4 (Turner, 2014). We considered associations to be significant when *p*-values were smaller than the Bonferroni threshold at *α* = 0.05 (*p* < 1.1e-07). False discovery rate (FDR) thresholds of 5% were determined using the *p.adjust* function in the stat package in R. We explored the genomic regions containing significantly associated loci using the “closest” command in bedtools v. 2.29.0 (Quinlan & Hall, 2010). Regions in the genome spanning the most significant associations were investigated for linkage disequilibrium patterns. We calculated the linkage disequilibrium *r*^2^ between marker pairs using the “option–hap-r2” in VCFtools v. 0.1.15 (Danecek *et al*., 2011) with “--ld-window-bp” of 10000. A heatmap was generated based on the *r*^2^ values with the R package *LDheatmap* v 0.99-7 (Shin *et al*., 2006).

### De novo genome assemblies, TE annotation, synteny analyses

We analyzed the locus surrounding the genes *Zt09_1_01590* and *Zt09_1_01591* in multiple completely assembled genomes of isolates collected in Switzerland, the United States, Australia and Israel covering the global distribution range of the pathogen (Badet et al., 2020). For synteny plots, the available repeat-masked chromosome-scale assemblies were analyzed using pairwise BLASTN. Information on BLAST hits among homologous chromosomes was visualized in R using the *genoplotR* package (Guy *et al*., 2010). We analyzed signatures of repeat induced point mutations (RIP) using The RIPper online tool available at https://theripper.hawk.rocks/ (van Wyk *et al*., 2019).

To analyze sequence polymorphism at the locus, we used draft genome assemblies of 432 isolates from previously analyzed field populations in the United States, Switzerland, Israel and Australia (Hartmann *et al*., 2017; Oggenfuss *et al*., 2020). Illumina short read data was obtained from the NCBI Sequence Read Archive under the BioProject PRJNA327615 (Hartmann *et al*., 2017) and PRJNA596434 (Oggenfuss *et al*., 2020). We used SPAdes version 3.14.0 to produce draft assemblies for each isolate (Bankevich *et al*., 2012). We ran the tool with the following settings: -k 21,33,55,75,95 --careful. De novo assemblies were annotated for TEs using the TE consensus sequences (https://github.com/crolllab/datasets) generated for the species (Badet *et al*., 2020). Consensus sequences were previously manually curated and renamed based on the three-letter classification system (Wicker *et al*., 2007; Bao *et al*., 2015). The curated consensus sequences were used for annotation of each individual de novo assembly using RepeatMasker version 4.0.8 with a cut-off value set to 250 (Smit & Hubley, 2015), ignoring simple repeats and low complexity regions. Further filtering of the TE annotation included: (1) removal of element annotations shorter than 100 bp, (2) merging of identical adjacent TE families overlapping by more than 100 bp, (3) renaming of overlapping TE families overlapping by more than 100 bp as nested insertions, and (4) grouping of interrupted elements separated by less than 200 bp into a single element using a minimal distance between start and end positions.

## Declarations

### Ethics approval and consent to participate

not applicable

### Consent for publication

not applicable

### Availability of data and materials

Illumina short reads were retrieved from the NCBI Sequence Read Archive (BioProject accessions PRJNA327615, PRJNA596434 and PRJNA650267) accessible from https://www.ncbi.nlm.nih.gov/sra. All other data are reported in Supplementary Information.

### Authors’ contributions

NKS and DC conceived the study, NKS and TB performed analyses, LA provided datasets, NKS and DC wrote the manuscript with input from all co-authors.

## Acknowledgements

We are grateful for advice and critical feedback on a previous version of this manuscript from Ursula Oggenfuss and Guido Puccetti. Seeds for the experiments were kindly provided by DSP Delley Inc.

