## Supplementary Figure for "Rapid sequence evolution driven by transposable elements at a virulence locus in a fungal wheat pathogen"

### Supplementary Figures

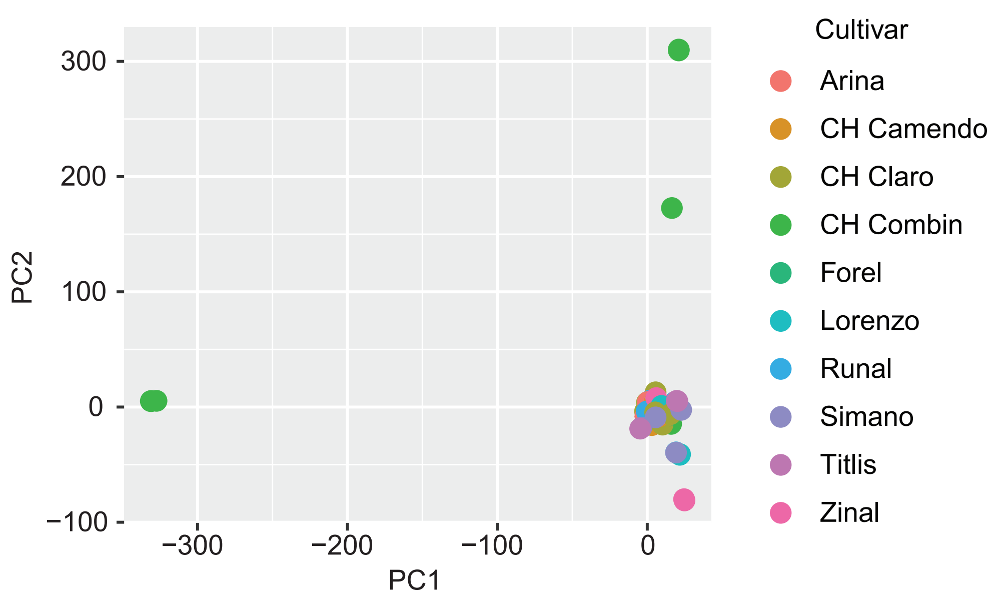


**Supplementary Figure 1 :** The first two principal components (PC) from a PC analysis of 788’313 genome-wide SNPs. Isolates are color-coded by the cultivar of the origin.

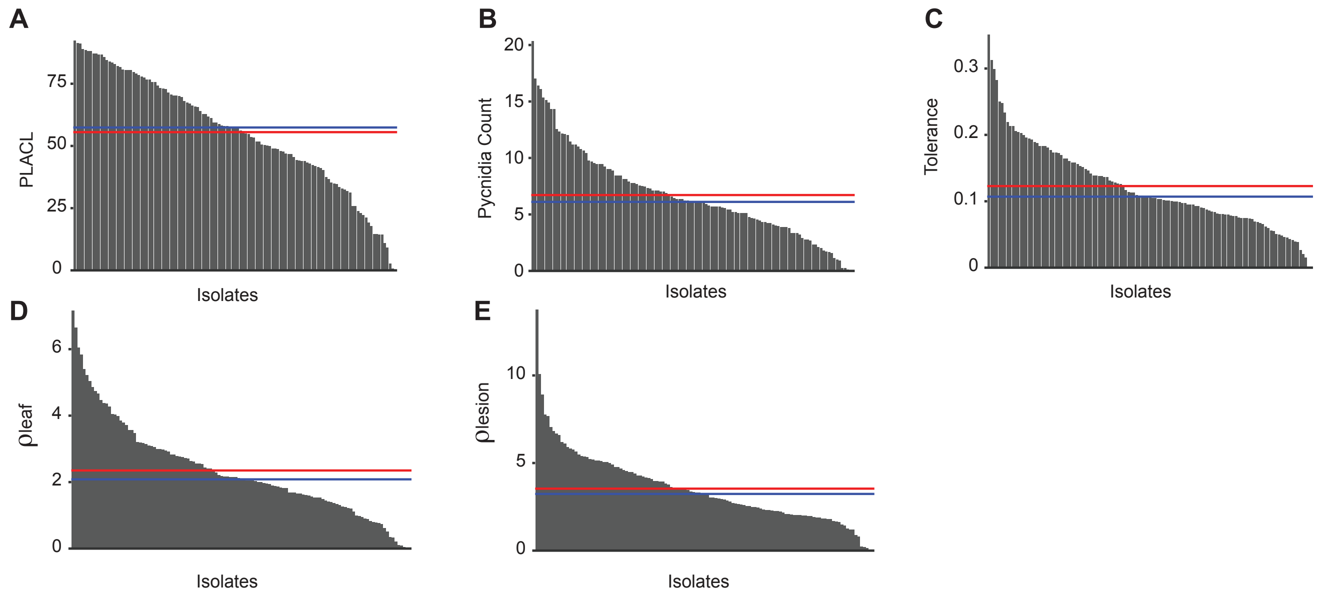


**Supplementary Figure 2 :** Phenotypic trait values. Red and blue lines represent the mean and median, respectively. A) The percentage of leaf area covered by lesion (PLACL). B) pycnidia count. C) Tolerance expressed as the pycnidia count divided by PLACL. D) $\rho_{leaf}$ is the pycnidia count per $cm^2$ of leaf area and E) $\rho_{lesion}$ is defined as the total number of pycnidia divided by per $cm^2$ lesion area.

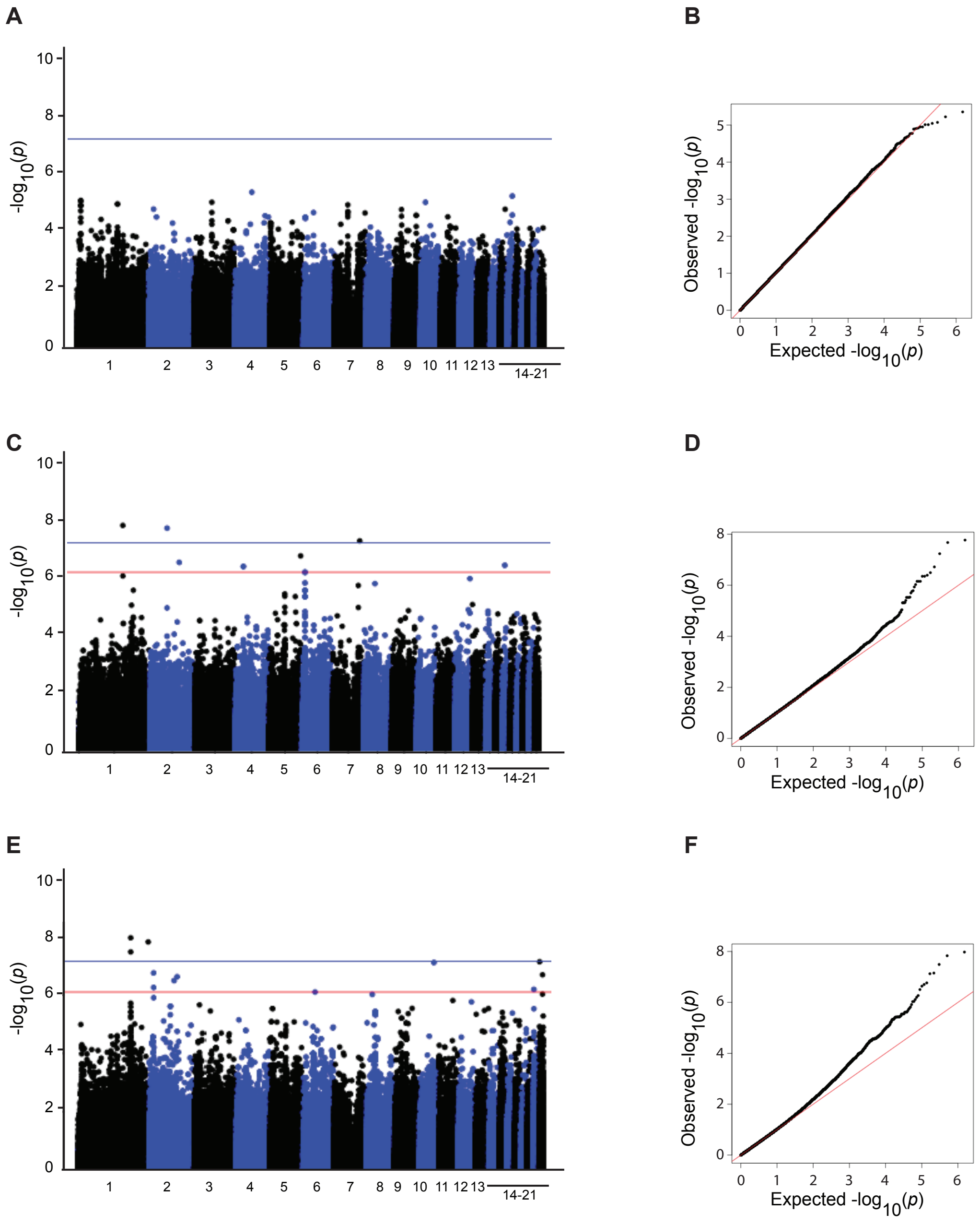


**Supplementary Figure 3 :** Manhattan and QQ-plot representing of the genome-wide association mapping analyses. The blue line corresponds to the Bonferroni threshod (alpha = 0.05) and the red line corresponds to the 5% FDR. A-B) Percentage of leaf area covered by lesions (PLACL), C-D) tolerance E-F) $\rho_{lesion}$

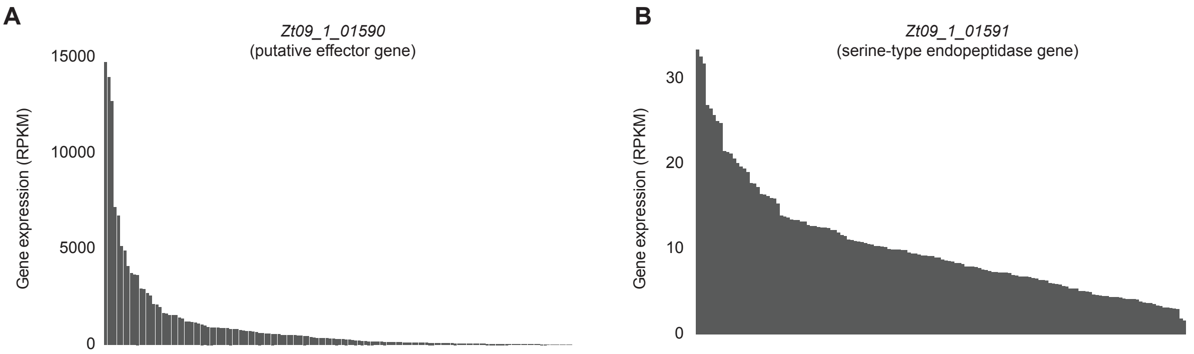


**Supplementary Figure 4 :** Gene expression in reads per kilobase of transcript, per million mapped reads (RPKM) for genes closest to the top-associated SNP. A) Putative effector gene (*Zt09_1_01590*) and B) serine-type endopeptidase gene (Zt09_1_01591).

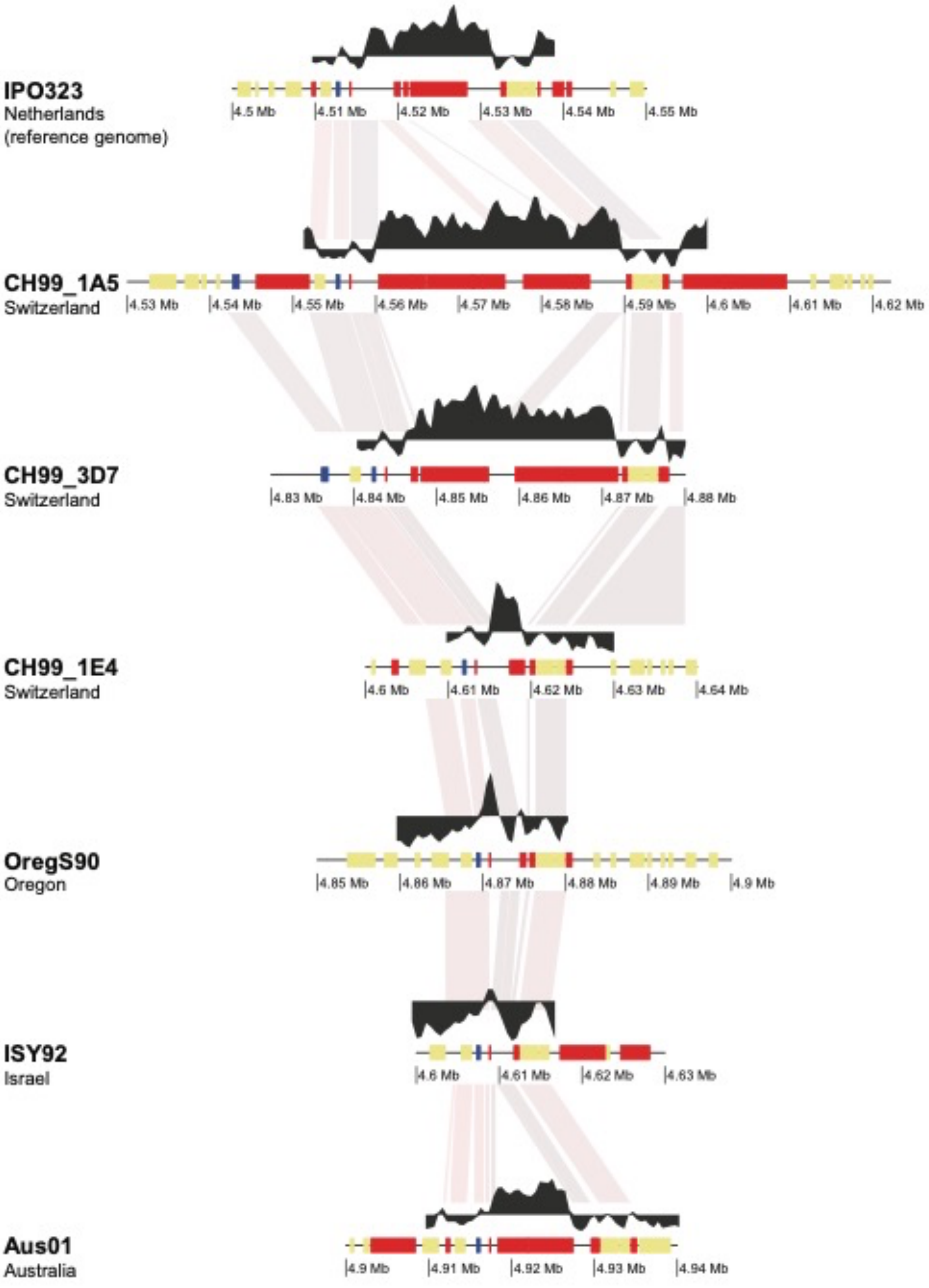


**Supplementary Figure 5 :** The Large RIP Affected Regions (LRARs) composite index was calculated using The RIPer tool (van Wyk et al., 2019) and shown in black. The region flanked by the genes *Zt09_1_01590* and *Zt09_1_01591* is shown in complete genome assemblies of seven isolates from global population.

### Supplementary Tables

(see separate Excel file)

**Supplementary Table S1:** Phenotypic trait values used for GWAS. The percentage of leaf area covered by lesion (PLACL); $\rho_{leaf}$ is the pycnidia count per $cm^2$ of leaf area; $\rho_{lesion}$ is defined as the total number of pycnidia divided by per $cm^2$ lesion area. Tolerance is expressed as the pycnidia count divided by PLACL

**Supplementary Table S2:** Groups of clonal genotypes identified in the GWAS population with information about the collection time point and cultivar of origin. The clonal genotype columns provides a unique identifier. See Singh *et al.* (2020) for more detailed analyses.

**Supplementary Table S3:** List of significantly associated SNPs above 5% FDR for pycnidia counts .

**Supplementary Table S4:** Number of isolates from populations on different continents analyzed for transposable element content. Total assembled genomes and total of isolates per population where a scaffold was retrieved containing both genes *Zt09_1_01590* and *Zt09_1_01591*.
